# High-content morphological profiling by Cell Painting in 3D spheroids

**DOI:** 10.1101/2025.02.05.636642

**Authors:** Christa Ringers, David Holmberg, Åsmund Flobak, Polina Georgieva, Malin Jarvius, Martin Johansson, Anders Larsson, Dan Rosén, Brinton Seashore–Ludlow, Torkild Visnes, Jordi Carreras Puigvert, Ola Spjuth

## Abstract

Cell Painting is a popular assay for morphological profiling of multi-labeled 2D monolayer cell cultures used in a wide range of applications. Culturing cells in 3D has potential for higher physiological relevance, such as when studying effects of perturbations. Robust and scalable 3D models can be challenging to characterize through imaging – particularly because light has difficulty penetrating cell multilayers. We introduce a scalable method where the Cell Painting assay is combined with tissue-clearing and applied to 3D spheroids generated in a ULA microplate format. Multi-channel images are acquired using confocal microscopy, and cells can be segmented inside those spheroids allowing for relevant morphological features to be extracted. Our end-to-end analysis pipeline comprises cell segmentation, morphological feature extraction, and between-spheroids and within-spheroid normalization. We demonstrate the method using spheroids cultured from two colorectal cancer cell lines and successfully detect distinct phenotypic changes upon compound treatments, on both spheroid-level using maximum intensity projections and on single cell-level. We show that drugs group by mechanism of action, with biologically relevant clusters especially evident with single-cell data. Finally, we contrast our method to results from 2D Cell Painting and discover a different pattern in DNA damaging drugs in HCT116 colorectal cancer cells. This work lays the foundation for multi-channel image-based screening in 3D spheroids.

## Introduction

Phenotypic profiling and cellular assays play important roles in early drug discovery, offering valuable insights that have helped uncover new disease mechanisms (Swinney & Anthony, 2011; Vincent et al., 2022). One approach is morphological profiling, where a large number of quantitative metrics (features) —such as cell size, shape, and texture—can be extracted from individual cells within images, creating a unique fingerprint for every given condition (Caicedo et al., 2017; Zhang et al., 2023). This shift towards large scale morphological profiling or phenomics allows for the exploration of less-characterized phenotypes, thereby facilitating the discovery of novel disease biomarkers (Houle et al., 2010; Kelley et al., 2023; Tegtmeyer et al., 2024).

Cell Painting is a widely adopted standardized assay for morphological profiling of 2D monolayer cell cultures. By staining distinct cellular compartments and imaging in multiple channels, it enables the extraction of vast amounts of phenotypic information. This unbiased approach is powerful for predicting mechanisms of action (Rose et al., 2018; Seal et al., 2024), identifying off-target effects (Schneidewind et al., 2020), and detecting phenotypes of environmental toxicants (Nyffeler et al., 2020; Rietdijk, Aggarwal, et al., 2022). Moreover, a growing variety of open-source software (Stirling et al., 2021; G. Way et al., n.d.) and publicly available datasets (Chandrasekaran et al., 2023) enhance the ease of conducting and comparing Cell Painting assays. Currently, the Cell Painting assay is primarily suited for 2D monolayers even though certain cellular functions—especially those that occur in three dimensions— remain challenging to capture in these types of models (Abbas et al., 2023; Däster et al., 2016; Riedl et al., 2017).

Three-dimensional (3D) cell cultures exhibit distinct biological characteristics compared to their 2D counterparts, where cells can engage in complex interactions with neighboring cells and, depending on the specific model, the extracellular matrix (ECM) (Nederman et al., 1984; Roskelley et al., 1994; Saraswathibhatla et al., 2023; Valdoz et al., 2021). The rigid 2D surface of a flask, on the other hand, directs cells to form large focal adhesions instead of making these intricate 3D interactions (Kanchanawong et al., 2010; Paszek et al., 2005; Saraswathibhatla et al., 2023) often leading to the alignment of cytoskeletal actin filaments into stress fibers (Gupta et al., 2015) – a phenotype not observed in 3D cultures (Saraswathibhatla et al., 2023). These differences in culturing methods do not only influence cell morphology but could also drive changes in cellular signaling (Riedl et al., 2017), such as promoting cell proliferation and impairing differentiation in 2D (Dupont et al., 2011; Paszek et al., 2005), while reducing proliferation (Pacheco-Marín et al., 2016; Pavel et al., 2018; Ramírez-Cuéllar et al., 2024) or diffusion of oxygen, nutrients, and waste in 3D (Hirschhaeuser et al., 2010; Pacheco-Marín et al., 2016). In addition, interactions with the 3D environment - shaping both a cell’s morphology and internal signaling - could alter their drug response, making morphological profiling experiments on 3D models increasingly relevant.

While considerable advances have been made towards scalable profiling in 3D (Betge et al., 2022; Huang et al., 2024; Lukonin et al., 2020; McCarty et al., 2023; Mertens et al., 2023; Sirenko et al., 2022), current methods do not capture single-cell level phenotypic variations. Individual images are usually condensed into Z-projections, effectively removing single-cell or lumen information, from which features are then extracted for further analysis (Booij et al., 2019; Shihavuddin et al., 2017). Light scattering in samples more than a few cell layers thick poses a major hurdle to single-cell 3D imaging as it complicates downstream cell segmentation and feature analysis (Egner & Hell, 2005; Silvestri et al., 2016). Recent advances using spinning disk confocal microscopy combined with either tissue-clearing or high Numerical Aperture (NA) objectives show that high-throughput detailed 3D imaging is possible (Boutin et al., 2018; Cutrona & Simpson, 2019). Nevertheless, its application in automated and multiplexed quantitative morphological profiling assays such as Cell Painting remains unchartered.

Here we introduce a scalable method for Cell Painting in 3D spheroids. We show that in combination with tissue-clearing, the protocol can be effectively applied to 3D spheroids formed in Ultra Low Attachment (ULA) plates. Using spinning disk confocal microscopy, we could acquire multi-channel images, enabling segmentation of individual cells frrom within the spheroids. Altogether, this work establishes a framework for multi-channel image-based phenotypic exploration in 3D spheroids.

## Results

### A Cell Painting workflow for 3D spheroids

The original Cell Painting assay was developed for single-cell morphological profiling of adherent cells in 2D. The assay, extensively reviewed and used in numerous applications (Caicedo et al., 2017; Nyffeler et al., 2020; Rietdijk, Tampere, et al., 2022; Rose et al., 2018; Schneidewind et al., 2020; Seal et al., 2024), involves the use of established dyes such as Hoechst, Syto14, Concanavalin A, Wheat Germ Agglutinin (WGA), Phalloidin, and MitoTracker to multiplex the visualization of subcellular organelles and structures (Bray et al., 2016; Caicedo et al., 2017). Here, we present an adaptation of the classical 2D Cell Painting assay for 3D spheroids, providing an end-to-end workflow from cell seeding to data processing (Figure 1).

**Figure 1:**
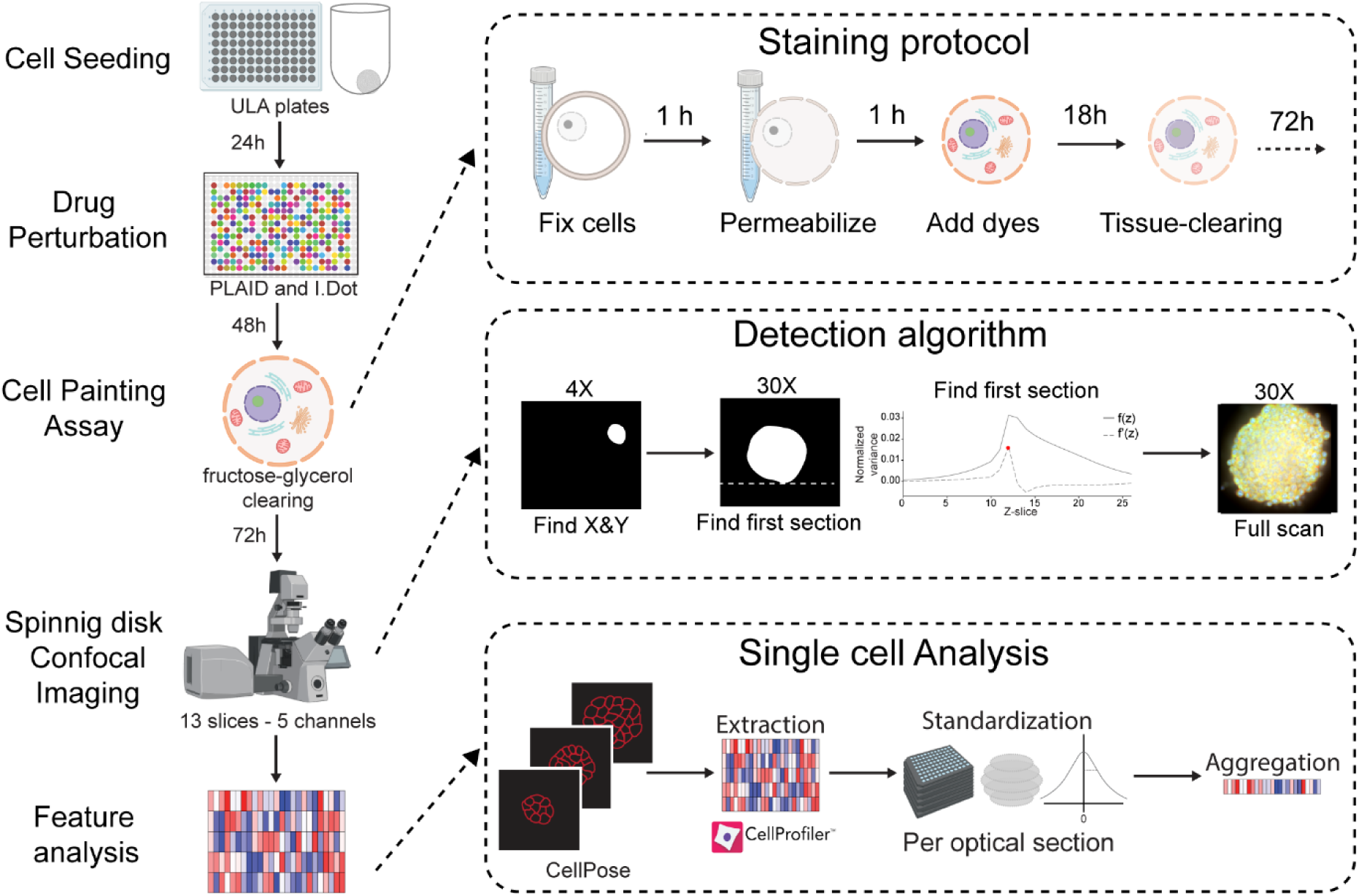
A workflow for Cell Painting of 3D spheroids. Schematic of the Cell Painting assay adapted to morphological profiling of spheroids (left). Cells are cultured in ultra-low attachment (ULA) plates (liquid-overlay) to form uniform spheroids. After 24h, spheroids are exposed to a drug library in a plate layout designed with PLAID. Spheroids are then subjected to the Cell Painting assay, followed by spinning disk confocal imaging and feature processing. Note the longer incubations and the fructose-glycerol clearing step. The staining protocol for spheroids in 3D. Fixation of cells (1h) is followed by permeabilization (1h), dye incubation (18h) and finally tissue-clearing by a fructose-glycerol mixture (72h; top-right). The spheroid detection starts with a scan at 4X to find the X-and-Y position, followed by a Z-scan to find the first optical section of each spheroid, and finalized by multichannel high detail imaging of the spheroid. Each spheroid is imaged across thirteen optical sections that are 5µm apart (middle-right). The analysis pipeline consists of cell and nuclei detection by CellPose, feature extraction by CellProfiler, per-section standardization and finally profile aggregation to collect per-well morphological profiles (bottom-right).

We used ULA plates (Sirenko et al., 2022) to produce uniform spheroids, simplifying cell seeding and enabling automation to improve assay throughput and robustness. To improve dye penetration, we extended the incubation time to 18 hours (Figure 1).

Imaging 3D dense spheroids is challenging due to light scattering (Egner & Hell, 2005; Helmchen & Denk, 2005; Ntziachristos, 2010; Richardson & Lichtman, 2015; Silvestri et al., 2016). To correct for this, we combine tissue-clearing and optical sectioning.

Tissue-clearing is a technique that homogenizes the imaging medium with the cellular and molecular components in a sample, effectively matching their refractive index (Silvestri et al., 2016). Initially developed to facilitate the microscopic visualisation of large *ex-vivo* organs in higher detail, the technique can now be successfully applied to spheroids and organoids for sub-cellular imaging (Dekkers et al., 2019; Diosdi et al., 2021). In our case, we opted for an aqueous clearing solution that achieves a high refractive index by mixing fructose and glycerol. This mixture, which is suitable for spheroids, balancing ease of preparation while providing good imaging results, and has been described previously (in similar or identical form) in work by others (Ariel, 2017; Dekkers et al., 2019; Diosdi et al., 2021; Nürnberg et al., 2020).

Optical sectioning with imaging techniques such as confocal or light-sheet microscopy, has historically and widely been used for visualising complex 3D biological structures (Keller & Dodt, 2012; Elliott, 2020; Boutin et al., 2018; McCarty et al., 2023; Mogollon et al., 2024). We took advantage of optical sectioning by spinning disk confocal to successfully image individual cells within cleared spheroids. The combination of tissue-clearing and optical sectioning enabled both the stains to diffuse to deeper layers of the spheroid and their visualization compared to non-cleared samples (Figure 2A-C, Supplementary 2A-B). Further building on this, and to preserve the assay’s scalability, we implemented automated imaging: a custom on-the-fly detection algorithm determined each spheroid’s initial Z position by analyzing focus changes instead of intensity across Z. In short, the algorithm involved low-magnification X-Y positioning (4x), high-magnification Z-scanning (20X), and lastly multi-channel Z-stack acquisition (Figure 1). This process ensured consistent Z position imaging across spheroids (Figure 2D), enabling homogeneous and stable image acquisition as well as carefully depth-matched and annotated sections.

**Figure 2:**
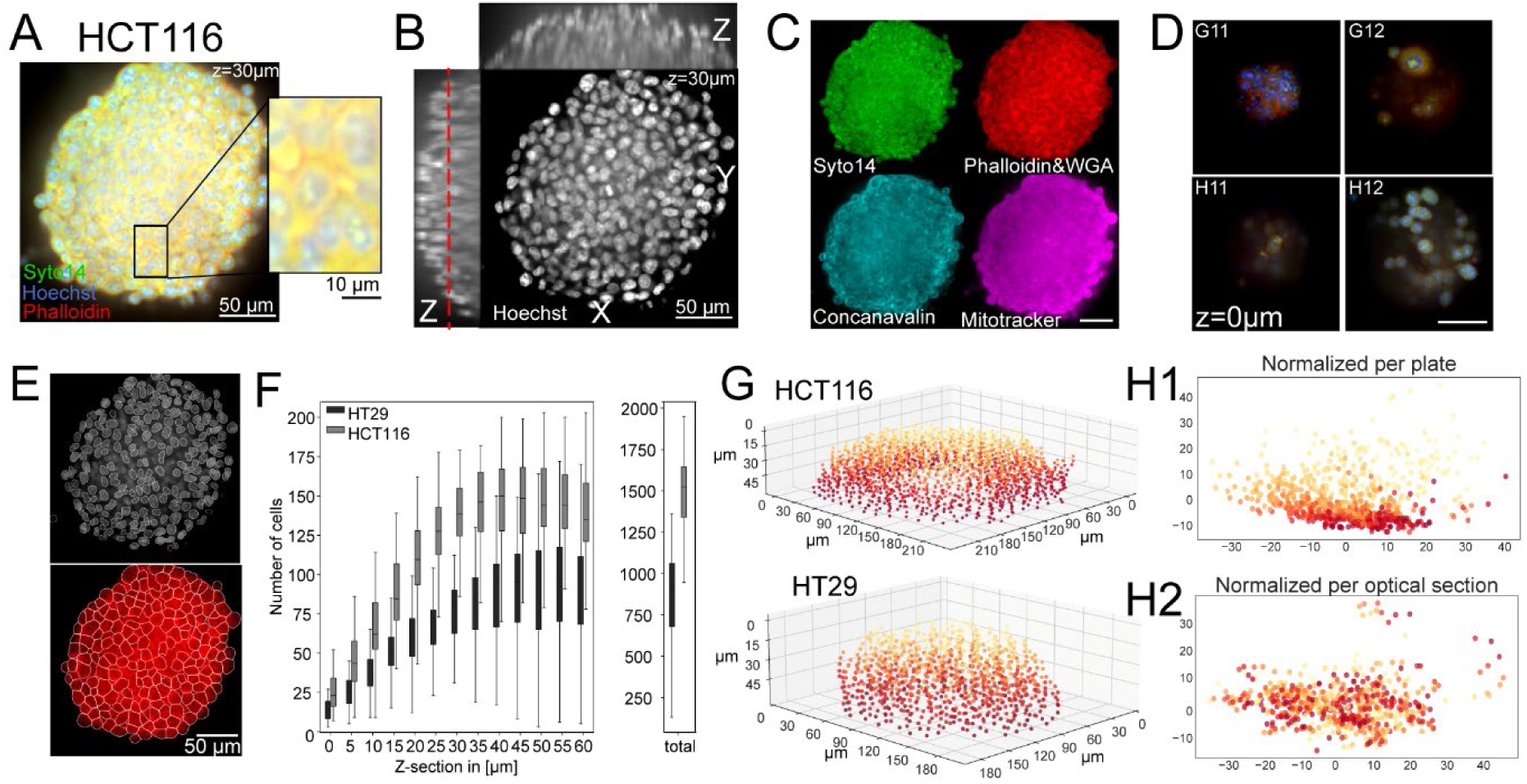
Towards reliable single-cell morphological profiles from cleared 3D spheroids. (A) Composite image of a HCT116 spheroid at 30µm depth depicting RNA and cytoplasm with (Syto14; green), nuclei with (Hoechst; blue) and cytoskeleton plus golgi (Phalloidin+WGA; red). Individual cells are seen in the inset. (B) Displayed are orthogonal views of the same spheroid in the Hoechst channel. These views consist of 12 made-isotropic slices (ranging from 0 to 60µm, with 5µm spacing). A dashed red line indicates the position of A (left). (C) A montage of individual channels: the cytoskeleton and Golgi are stained with phalloidin and WGA (red), mitochondria are stained with MitoTracker (magenta), nuclei are stained with HOECHST (blue), ER is stained with Concanavalin A (cyan), cytoplasm and RNA are stained with Syto14 (green). (D) Representative images of detected spheroids. They represent the first optical sections as found by the detection algorithm. (E) An example of nuclei (top) and cell segmentation (bottom) by CellPose. Objects are outlined in white. (F) A summary of the number of detected cells per section for the cell lines HCT116 and HT29. (G) The center positions of detected cells in HCT116 (top) and HT29 (bottom) spheroids exposed to 0.1% dmso. (H) A PCA of dmso morphological profiles section-aggregated before (top) and after (bottom) per-section standardization. Spheroid depth is color-coded in G-H. Scale bars are 50µm, unless indicated otherwise.

For cell segmentation, we used CellPose - a tunable deep learning-based model (Pachitariu & Stringer, 2022) - to automatically segment cells from each acquired image section (Figure 1). We reasoned that a 2D approach should still allow us to capture morphologies that are representative of their 3D biological structure. In this way, we could resolve single cells inside spheroids (Figure 2E), enabling the average per spheroid segmentation of 1425 cells (mean ± SD 352) in HCT116 and 854 cells (mean ± SD 274) in HT29 (Figure 3F-G), from which we could then extract >2000 morphological features using Cell Profiler.

**Figure 3:**
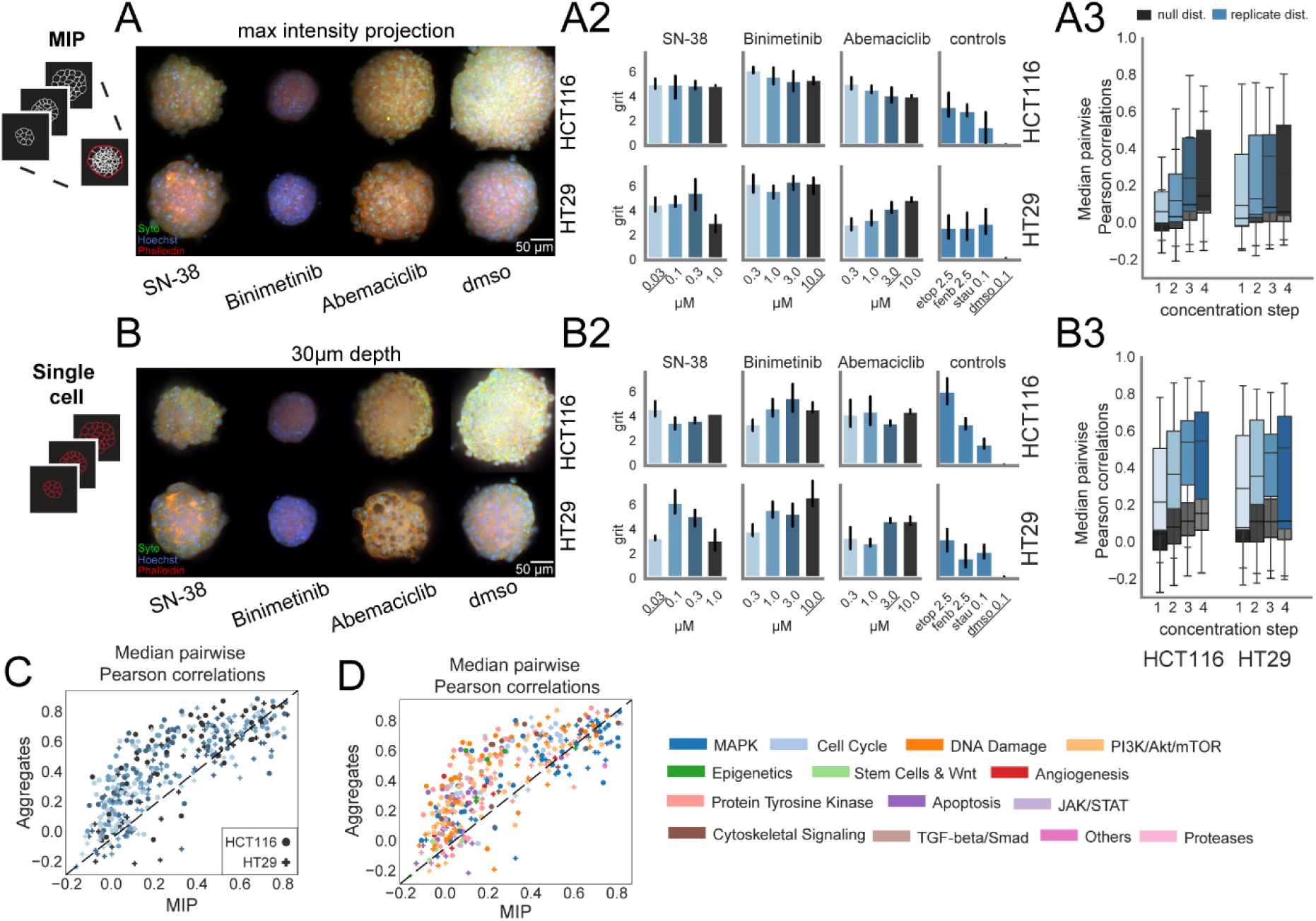
Comparing the quality of single-cell level profiles to spheroid-wide morphological profiles. (A) Representative examples of HCT116 (top) and HT29 (bottom) MIPs for the compounds SN-38 (0.03 uM), binimetinib (10 µM), abemaciclib (3.0 uM), and dmso (0.1%), as well as for (B) a single section at 30µm depth. Syto (RNA) is shown in green, Hoechst (nuclei) is shown in blue, and Mitotracker (mitochondria) is shown in red. (A2-B2) Grit scores measuring phenotypic effect are shown for MIP (A2) and aggregated (B2) profiles. Error bars indicate bootstrapped 95% confidence intervals. (A3-B3) Null and median pairwise replicate correlations distributions across dose steps for both HCT116 and HT29 cell lines. Null distributions are in grays and replicate in blues. (C) Comparison of median pairwise replicate correlations of MIP and aggregates profiles in both HCT116 and HT29 cell lines. Data points are either color-coded for (C1) dose, or (C2) pathway.

Despite these optimizations, a depth-dependent decrease in intensity and contrast was still noticeable (Figure 2B, Supplementary 2B), contributing to signal degradation observed in section-aggregated single-cell morphological profiles (Figure 2H1). However, by having carefully matched and annotated spheroid sections, we could address this issue. We normalized optical sections – equivalent sections across spheroids in a multi-well plate - against negative controls in the same section (Figure 1). This normalization strategy effectively corrected the signal degradation (Figure 2H2), enabling the generation of morphological profiles for downstream analysis (Caicedo et al., 2017).

While some aspects of the workflow depend on specific imaging systems, such as the spinning-disk, type of objective and its lasers, the underlying principles—tissue-clearing, automated segmentation, and normalization—are adaptable to other platforms. These advancements maintain the efficiency and scalability of the original Cell Painting assay while enabling robust 3D profiling in spheroids.

### Single-cell level profiling captures phenotypic changes with greater resolution compared to spheroid-wide profiling (MIP)

To evaluate our 3D Cell Painting workflow, we profiled two colorectal cancer cell lines (HCT116 and HT29) treated with 52 clinically relevant drugs and pre-clinical compounds, from now on referred to as drugs (Supplementary Figure 3A; Supplementary Table 1). We also included the phenotypic reference compounds etoposide and fenbendazole (Willis et al., 2020). In short, cells were seeded in ULA plates 24 hours before exposure to allow for spheroid formation. We designed effective plate layouts using the constraint programming tool PLAID (Francisco Rodríguez et al., 2022). After 48 hours of drug exposure, we performed our 3D Cell Painting workflow followed by fructose-glycerol clearing for 72 hours, making the entire protocol span one week. Using this approach, we successfully stained and imaged spheroids in 1607 wells. Yet, some treatments might have increased the fragility of the spheroids, resulting in them being washed away during the staining process. We then continued to profile those spheroids using CellPose and CellProfiler (Figure 1). However, 3D spheroid feature processing techniques differ from 2D. Single-cell data from 2D Cell Painting experiments are commonly aggregated into median cell features (Caicedo et al., 2017; Ljosa et al., 2013). 3D morphological profiling experiments often rely on Z-projections as a widely used method for feature extraction (Betge et al., 2022; Booij et al., 2019; Huang et al., 2024; Lukonin et al., 2020; Mertens et al., 2023; Sirenko et al., 2022). One such technique is the maximum intensity projection (MIP) – where a pixel with the highest intensity across a Z-stack is selected (Figure 3A). Yet, this way images may lose relevant single-cell information and contain artifacts - such as objects appearing closer than they really are due to differences in Z positioning (Booij et al., 2019; Shihavuddin et al., 2017).

We compared the morphological profiles of features extracted both at the spheroid-wide level (using MIP; Figure 3A) and the single-cell level (using aggregates; Figure 3B). Several drug treatments resulted in visible phenotypic changes. For example, spheroids exposed to binimetinib are smaller and appear to have small cytoplasmic compartments; spheroids exposed to SN-38 seem to have fewer cells that are larger; and spheroids exposed to abemaciclib could have formed lysosome-derived vacuoles (Hino et al., 2020)(Figure 3A1- B1). We then turned to the actual morphological profiles. To measure the strength of phenotypic changes we calculated grit scores for both the MIPs and single-cell aggregates, indicating how much more similar a drug is to its replicates than to the negative controls (Trapotsi et al., 2022; G. Way et al., 2020/2023, 2020/2024). These scores are directly interpretable where a grit of 1.96 represents the cutoff for the 95% confidence interval, serving as a threshold to identify meaningful phenotypic shifts. We compared the number of drugs exceeding this threshold between MIP profiles (Figure 3A2) and single-cell aggregate profiles (Figure3B2). We find that 38 (grit: µ=2.01±1.78; MIP) and 37 (grit: µ=1.86±1.95; aggr.) drugs for HCT116, and 33 (grit: µ=1.92±1.95; MIP) and 32 (grit: µ=2.10±2.26; aggr.) drugs for HT29 had a grit score over 1.96. The detection of phenotypic shifts is thus comparable for the two streams of preprocessing, indicating that morphological profiles of perturbed spheroids can be successfully distinguished from negative controls in both MIPs and single-cell aggregates.

To evaluate the resolution of morphological profiles and the reproducibility between technical replicates of the assay we used the ‘*percent replicating*’ metric, which is based on median Pearson replicate correlations and captures the proportion of profile replicates that are more similar to one another than one would expect by chance – specifically compared to a null distribution generated by randomly sampling non-replicates (Cimini et al., 2022; G. P. Way et al., 2022). We find that the percentage increases with concentration (Figure 3A3-B3), is better for the HCT116 cell line (Figure 3A3-B3), and slightly lower than previously published values – although values may fluctuate depending on compounds included in an experiment (Cimini et al., 2022; G. P. Way et al., 2022). When comparing *percent replicating* between spheroid-wide (MIP) or aggregated single cell, we indeed observe higher values when extracting features from single cells (Figure 3C-D). This effect does not seem to be carried by cell line, concentration (Figure 3C), or pathway (Figure 3D), This indicates that while both MIPs and aggregate profiles are different and distinguishable from negative controls, it is the single-cell aggregate profiles that can be discriminated from each other. In other words, those profiles have a better resolution.

### Single-cell morphological profiles facilitate the detection of biologically relevant clusters

Morphological profiling via Cell Painting has been used to profile compounds based on their bioactivity, assess drug safety, and predict compound mechanism of action (MoA)(Bianchi et al., 2024; Garcia de Lomana et al., 2023; Seal et al., 2021; G. P. Way et al., 2022). A central aspect of these tasks is leveraging morphological profiles to identify phenotypically similar clusters of compounds. To evaluate the potential of our MIP and single-cell aggregated profiles to discriminate drug clusters from perturbed 3D spheroids, we first explored Uniform Manifold Approximation and Projection (UMAP) (McInnes et al., 2020) – a unsupervised method commonly used for downstream analysis of biological features (Cao et al., 2019). We included class labels – here pathways - in the clustering algorithm to account for data variability. This allowed us to identify separate clusters per pathway in the UMAPs for MIP and aggregated profiles in HCT116 and HT29 cell lines (Figure 4A1-B1; Supplementary 4A1-B1).

**Figure 4:**
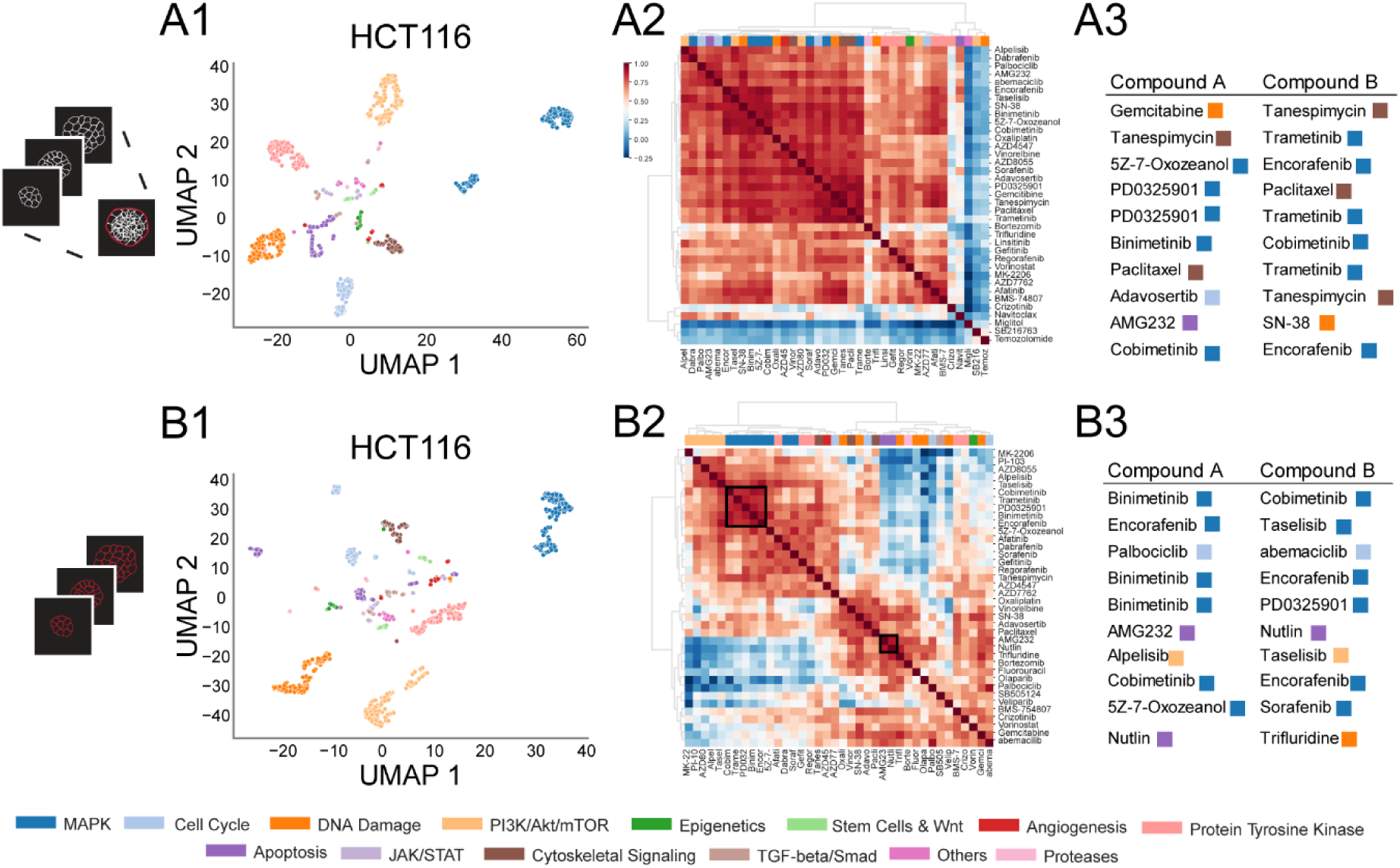
Single-cell morphological profiles facilitate the detection of biologically relevant clusters. (1) Semi-supervised UMAPs of MIP (A1) and aggregated (B1) morphological profiles. (2) Heatmap displaying hierarchical clusters of cosine similarities for MIP (A2) and aggregated (B2) profiles. The ward algorithm was used to create the dendrogram. (3) Top 10 most similar compounds for MIP (A3) and aggregated (B3) profiles. In all plots, pathways are color-coded using the same colormap. All data is based on HCT116 spheroids.

To complement our UMAP analysis, we performed hierarchical clustering (using cosine similarity) to generate cluster maps, excluding any drug with grit scores less than 1.96. While we observe a single cluster in the spheroid-wide MIP profiles (HCT116; Figure 4A2), we observe distinct clusters in the single-cell aggregated profiles (HCT116; Figure 4B2); one cluster primarily includes kinase inhibitors (MEK, PI3K, and PTK), and the other contains various types of drugs including DNA damaging drugs.

Notably, we find the phenotype of two cell cycle inhibitors – abemaciclib and palbociclib, targeting CDK2/4, to be highly similar (cosine sim = 0.89) in both cell lines. We also find two MDM2 inhibitors (Koo et al., 2022), nutlin-3a and navtemadlin (AMG232) to be highly similar in HCT116 (cosine sim = 0.89; Figure 4A3), but not in HT29 (Supplementary Figure 4A3). Indeed, the HCT116 cell line is WT for TP53, and therefore expected to be more sensitive to MDM2 inhibition (Tovar et al., 2006). These results highlight the potential of these profiles to uncover biological relationships within our 3D dataset, offering an initial approach for understanding the intricate patterns underlying drug activity and cellular responses at the single-cell level.

### Morphological profiles reveal different responses to DNA damaging drugs in spheroids versus adherent cells

While viability screens have reported differences in responses between adherent (2D) or spheroid (3D) cultures (Folkesson et al., 2020; Tung et al., 2011; Vinci et al., 2012), differences in morphological profiles between these modalities remain largely unexplored.

We performed Cell Painting in 2D on HCT116 and HT29 cell lines and generated single cell aggregate morphological profiles for our 52 drug perturbations (Figure 5A; Supplementary 5A). To compare the assays’ ability to detect phenotypic shifts, we calculated grit scores, observing strong shifts for 2D perturbations (Figure 5B; Supplementary 5B). We find that 46 (grit: µ=3.90±2.65; 2D) and 37 (grit: µ=1.86±1.95; 3D aggr.) drugs for HCT116, and 47 (grit: µ=3.91±2.91; 2D) and 32 (grit: µ=2.10±2.26; 3D aggr.) drugs for HT29 had a grit score over 1.96. It is unclear why there are fewer phenotypic drug responses in 3D. It may reflect technical limitations of the 3D workflow but could also point towards a stronger resistance of 3D spheroids to drug perturbations.

**Figure 5:**
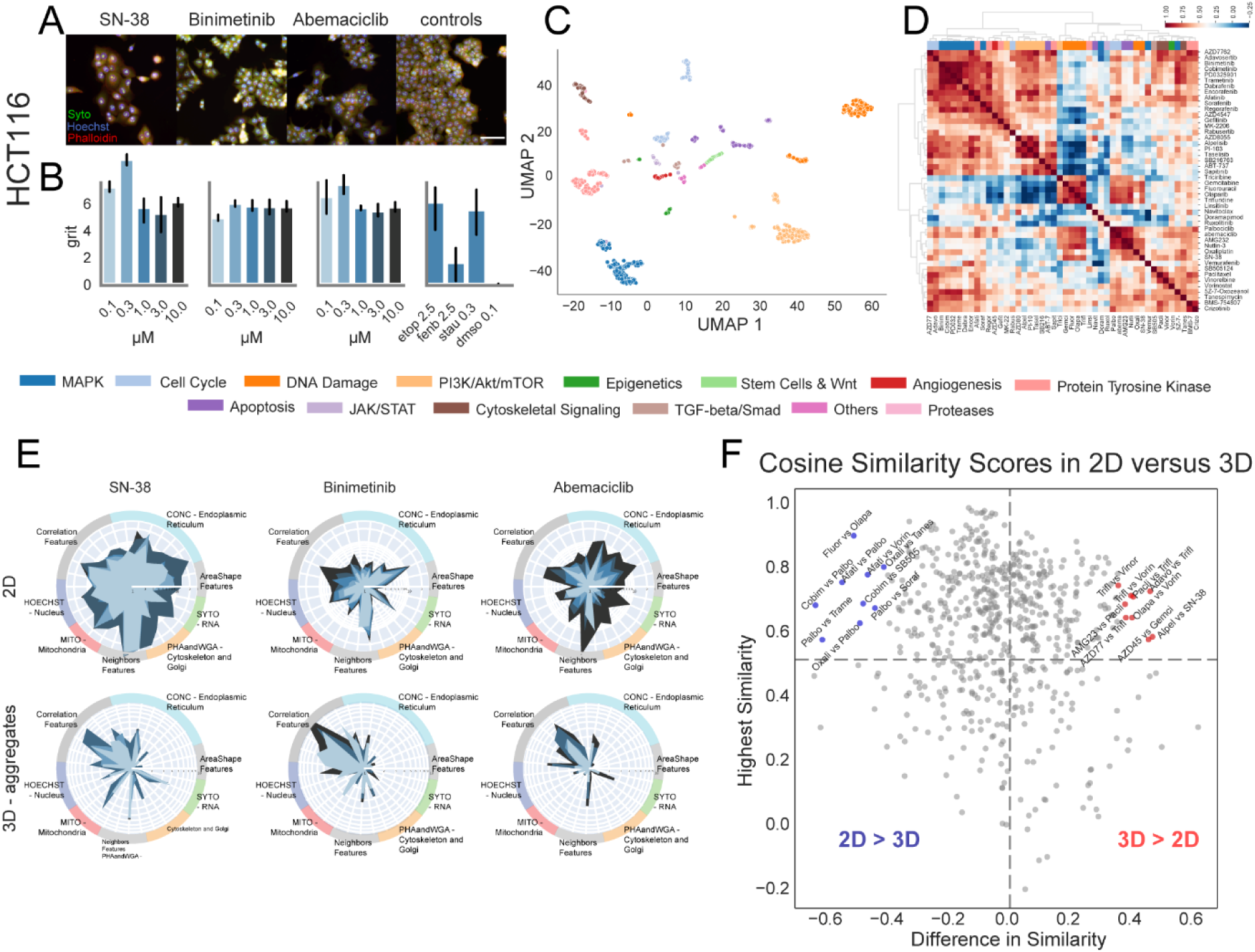
Morphological profiles reveal different responses to DNA damaging compounds in spheroids versus adherent cells. (A) Representative examples of HCT116 cultured in 2D for the drugs SN-38 (0.03 uM), binimetinib (10 µM), abemaciclib (3.0 uM), and dmso (0.1%). (B) Grit scores measuring phenotypic effect are shown for 2D aggregate feature profiles. Error bars indicate bootstrapped 95% confidence intervals. (C) Semi-supervised UMAPs of 2D morphological profiles. (D) Heatmap displaying hierarchical clusters of cosine similarities for 2D profiles. (E) Radar plots for the same drugs are displayed for 2D (top) and 3D aggregate (bottom) morphological profiles. Doses are color-coded according to the colors used in B. (F) Comparison of cosine scores between pairs of compounds in 2D and 3D. The x-axis shows the difference in similarity scores (3D – 2D), while the y-axis shows the highest similarity score across both modalities for each pair. We only included pairs with a cosine score of at least 0.5 in either 2D or 3D, and that exceeded the 95% confidence interval cutoff. All data is based on HCT116 cells. Scale bar is 250 µm.

Nevertheless, UMAP plots of 2D and 3D perturbations reveal comparable clusters, indicating that both modalities capture biologically relevant information (Figure 5C; Supplementary 5C).

We then performed cosine similarity clustering of perturbations in 2D. For HCT116, we could readily observe clusters of MAPK and PI3K/Akt/mTor in both modalities, but not the DNA damaging compound cluster, which was present in 2D only (Figure 4B2; Figure 5D), potentially suggesting a different mechanism of these drugs in 2D versus 3D. DNA damaging drugs have indeed been shown before to have a different effect on adherent and 3D cultures (Tung et al., 2011).

To further compare specific morphological profiles, we generated radar plots (Rietdijk, Aggarwal, et al., 2022) for the drugs SN-38, binimetinib and abemaciclib. These plots revealed individual patterns for each drug that become more pronounced with increasing concentrations (Figure 5E) yet seem more pronounced for correlation features than other types of features. We do not see this in the 2D radar plots. This modality-specific effect may result from signals captured from adjacent optical sections, which are slightly above or below the focal plane in 3D.

Recognizing the strength of morphological profiling in identifying similarities between perturbations (Seal et al., 2024), but also the limitations when it comes to directly comparing profiles between experimental conditions such as batches (Arevalo et al., 2024; Sypetkowski et al., 2023), cell resistant clones (Kelley et al., 2023), and imaging modalities, we instead focused on exploring how similarity patterns manifest between 2D and 3D.

To this end, we systematically compared cosine similarities across drugs treatments in 2D and 3D, emphasizing pairs with strong modality-specific variation, defined as those falling above the 95^th^ percentile of the observed differences (Figure 5F; Supplementary 5F). For instance, DNA-damaging drugs 5-Fluorouracil and Olaparib showed high similarity in 2D but not in 3D (Figure 5F), indicating a potential modality-dependent mechanism of action. Interestingly, neither drug induced measurable cell death in 2D or 3D, as determined by a metabolic activity assay (Folkesson et al., 2020). Despite this, both drugs elicited phenotypic responses, highlighting the use of morphological profiling in capturing functional readouts even in absence of measurable cytotoxicity. We observed differences in 2D versus 3D with respect to DNA damaging compounds, likely reflecting biological characteristics between these culture modalities. While morphological profiling can uncover important phenotypic variations, interpreting these differences may benefit from complementary approaches.

## Discussion

In this work, we introduced and demonstrated a new method and workflow for scalable Cell Painting in 3D. By combining fructose-glycerol clearing, CellPose segmentation, and per-section normalization to mitigate depth-related changes in image quality, our approach allowed for extracting single-cell features. We showed that segmentation of cells within spheroids is possible, and that their morphological fingerprints carry relevant biological information. As compared to features extracted from max intensity projections (MIP), these single-cell features provide better resolution, even though both types of features can distinguish spheroids treated with small molecules from those untreated. In addition, we identify differences in 3D profiles as compared to 2D, specifically with respect to DNA damaging compounds.

We anticipate that 3D Cell Painting will enable morphological profiling of culture models that are known to be difficult to capture in 2D, such as those exhibiting unique morphologies in 3D (Kenny et al., 2007; McCaffrey & Macara, 2011), such as liver culture models, which tend to flatten and lose some of their function in 2D (Bachmann et al., 2015; Kostadinova et al., 2013), or such as models that co-culture immune cells (Courau et al., 2019). Our method may also enable systematic exploration of spheroids from various tissue types, uncovering novel phenotypic biomarkers linked to 3D cellular complexity, as well as inspire the development of methods to perform single-cell morphological profiling in 3D cultures beyond spheroids.

While 2D cell cultures are very suitable for high-throughput experiments, 3D cultures remain less standardized, with diverse methods and terminology (Gunti et al., 2021; Langhans, 2018; Peirsman et al., 2021). Using ULA plates is one way to culture cells in 3D demonstrating compatibility with scalable morphological profiling. It would be interesting to extend our single cell profiling approach to more complex models that capture different aspects of *in vivo* biology, ultimately contributing to reducing the need for animal experiments (Bédard et al., 2020).

Even though our single-cel morphological profiling approach is effective, it also has certain limitations. We detected cells in each plane independently, treating them as 2D images. In this process, we likely sampled each cell more than once. It is possible to segment cells in 3D to avoid such overrepresentation (Diosdi et al., 2024; Mogollon et al., 2024; Moos et al., 2024; Weigert et al., 2020), along with the extraction of 3D features (McQuin et al., 2018). It remains to be tested how these morphological profiles capture biological information.

We normalized sections separately and aggregated profiles at the spheroid-wide level. This approach enabled a broad analysis, but it also collapsed single-cell information and implicitly assumed minimal biological variation with depth. However, given that spheroids often contain distinct spatial zones (Hirschhaeuser et al., 2010), future work distinguishing between technical and biological depth variations could further improve the accuracy of the method. Recently, several Deep Learning models have been published that can replace CellProfiler feature extraction (Kraus et al., 2024; Moshkov et al., 2024). Incorporating depth information into a deep learning feature extraction method that is imaging depth-invariant would be exciting, potentially enabling scalable, automated 3D single-cell analysis. Nevertheless, we showed how to implement and analyse a 3D Cell Painting protocol and uncovered differences in 2D versus 3D with respect to DNA damaging compounds.

Olaparib, a PARP inhibitor, traps PARP1 onto DNA, leading to replication stress and cell cycle arrest (Bryant et al., 2005; Farmer et al., 2005; Murai et al., 2012). In contrast, 5-Fluorouracil, a nucleobase analogue, disrupts DNA and RNA integrity, triggering apoptosis depending on the cellular context (Chen et al., 2024; Ewald et al., 2008; Longley et al., 2003; Pettersen et al., 2011). These drugs exhibited distinct responses in 3D but not in 2D cultures, potentially due to differences in proliferation and intracellular processing (Dupont et al., 2011; Paszek et al., 2005; Riedl et al., 2017). This highlights the benefit of 3D morphological profiling in uncovering modality-specific drug effects, particularly for DNA-damaging agents. Further studies are needed to clarify the underlying mechanisms driving these differences.

## Methods

### Cell lines

Two human colorectal cancer cell lines were used: HCT-116 (CVLC_0291) and HT-29 (CVLC_0320). Cell lines were routinely tested for mycoplasma using a luminescence-based Lonza Mycoplasma kit (Lonza™ LT07-418) and tested negative. In addition, cell lines were authenticated successfully by STR profiling (Eurofins genomics GmbH). Both cell lines were cultured in RPMI 1640 medium (Gibco 31870) supplemented with FBS (10%; Sigma F7524), Penicillin-Streptomycin (100U/ml; Gibco 15140-122), and GlutaMax (1X; Gibco A1286001). Cells were kept at 37 °C in a humidified atmosphere at 5% CO_2_ and passaged when confluency reached 80-90% using Trypsin-EDTA (0.25%; Gibco 15400054) for dissociation. Cells used in experiments never exceeded passage 10 after thawing.

### Clearing solution

Prior to the experiment, we prepared 100 ml of clearing solution by mixing 40 ml glycerol (40% vol/vol), 40 g of fructose (40%; wt/vol; 2.2202 M), and 32 ml of dH20 on a magnetic stirrer. This solution has a refractive index of 1.452-1.451 at RT as measured with a refractometer.

### Compound library and compound spotting

A clinical library encompassing 52 small molecules (Supplementary Table 1) was manually curated by relevance for (colorectal) cancer therapy and mechanism of action. Available drugs were sourced from SelleckChem as a Cherry Pick Library. The remaining drugs AMG232 (TA9H93ED7611; Sigma) and 5Z-7-OXOZEAENOL (O9890; Sigma) were purchased separately from Sigma-Aldrich. Five concentrations per compound were screened (100-fold dilutions). Compound libraries were stored in an airtight container at −20 °C before use. We included three phenotypic reference compounds with a known distinctive phenotype (Willis et al., 2020): Etoposide (E1383; Sigma), Fenbendazole (F5396; Sigma). Furthermore, Staurosporine (19-123; Sigma) was included as cytoxic control (Warchal et al., 2020), D-Sorbitol (S1876;) as negative control, and DMSO (D2438) as compound vehicle (purchased from Sigma Aldrich).

To reduce the impact of plate effects, we both excluded outer wells and designed effective microwell layouts using constraint programming with PLAID - Plate Layouts using Artificial Intelligence Design (Francisco Rodríguez et al., 2022). Specifically, we ran PLAID protocols through MiniZincIDE (Version 2.6.4), followed by a custom Jupyter Notebook to translate PLAID protocols to I.DOT protocols. We spotted drugs onto Corning® 384 well microplates (CLS 3656) using the I.DOT (I.DOT; Dispendix, Stuttgart, Germany) with the I.DOT PURE S Plate 100 (D16110021800). When plates were not used immediately, they were sealed and stored at −20 °C. Before use, we first dissolved the drugs in 25µl cell medium (4X). Afterwards we centrifuged (50RCF; 20s) and incubated the plates on a shaker at RT (20min). For 2D experiments, drugs were directly spotted onto PhenoPlate™ 384-well microplates (Revvity; 6057302).

### Cell seeding and compound treatment

We seeded cells using the ViaFlo 384 (Integra) in 30µl cell medium on 384-well ULA spheroid microplates (Corning; 3830). Both cell lines were seeded at 300cells/well (HCT116; p+10 / HT-29; p+8) in all wells. To support spheroid aggregation, we centrifuged (120RCF; 2min) and incubated the plates (24h; 37 °C; 5% CO_2_). Afterwards, we exposed the spheroids by transferring 10µl of drugs in cell medium using the ViaFlo to make a total of 40µl.

For 2D experiments, we first dispensed 20µl of cell medium in 384-well plates and incubated them at room temperature for 20 minutes, followed by dispensing 20µl cell mixture using the Biotek MultiFlo FX (HCT-116, 2600cells/well, p+8; HT-29, 2000cells/well, p+6). Finally, the plates were kept in an incubator (24h; 37 °C; 5% CO_2_).

### The Cell Painting assay

Following 48h compound treatment, we performed the Cell Painting Assay (Bray et al., 2016; Cimini et al., 2022) either in 2D or adapted to accommodate for 3D spheroids. For both experiments, we implemented wash steps with the Biotek 405 LS and dispense steps with the Biotek MultiFlo FX.

#### 3D protocol

We washed the cells once with 60µl 1X PBS (Gibco™ 18912014), incubated 4min, and aspirated leaving 20µl in the well. Then, we fixed with 60µl 4% PFA (Histolab; 02176) for 1h at room temperature. We washed twice (60µl, 5min wait, 1X PBS, leaving 20µl) and permeabilized with 60µl Triton X-100 0.2% (final well concentration; Thermo Scientific 85111) for 1h at room temperature, washed once (60µl, 1X PBS, leaving 10µl), and finally stained with 20µl stain mixture (see Supplementary table 2) over night (16-18h) at 4C. The next day, we washed twice (60µl, 2.5min wait, 1X PBS, leaving 10µl) and dispensed 70µl of clearing solution. We sealed and kept plates for 72h at 4°C prior to imaging.

#### 2D protocol

We washed the cells once with 80µl 1X PBS (leaving 20µl). Then, we dispensed 40µl MitoTracker (Invitrogen; M22426) in prewarmed 37°C FluoroBrite (Gibco™ A1896701) and incubated for 20 min (37°C, 5%CO2). Then, we washed the cells once (80µl, 1X PBS, leaving 20µl), fixed with 80µl 4% PFA for 20 min (RT), washed three times (80µl, 1X PBS, leaving 10µl), permeabilized with 80µl Triton X-100 0.1% (20min, RT), washed three times (80µl, 1X PBS, leaving 10µl), stained with 20µl stain mixture (Supplementary table 2; 20min, RT), and finally washed three times (80µl, 1X PBS, leaving 80µl). We sealed and kept plates at 4C prior to imaging.

### Image acquisition 3D

We imaged the plates in the Nikon ECLIPSE Ti2-E inverted microscope using a 20X objective (PLAN APO λD 20X OFN25 DIC N2, wd = 0.8mm, NA=0.8) combined with an additional 1.5X additional zoom, to create a total zoom of 30X (res=0.227 µm/pixel). The microscope was fitted with a Celesta light engine and 50µm spinning disk (spacing=250µm). Each channel was imaged with the filter settings and exposure times described in the Supplementary table 2. To detect the spheroids, we used a custom protocol in Nikon JOBS and GA3 (See Supplementary material). In summary, we scanned every well at 4X in the SYTO channel (widefield, binning=2x, exposure time=5ms, laser=0.42). By thresholding, we automatically estimated the position of each spheroid (threshold=1424). At each measured location, the entire spheroid was first scanned at 20X in SYTO (range=550µm, step=10µm, confocal=50µm, exposure time=50ms, laser=0.42). From this Zstack we would calculate the Z differential of the FocusScore - or normalized variance (Stirling et al., 2021; Sun et al., 2004). A higher Z-differential, or in other words a sharp change in Focus would indicate the start of a spheroid. At this measured Z location, we started imaging the spheroid in 5 channels (Supplementary table 2), with 5µm spacing across 12 optical sections. Images were saved as 16-bit ome.tiff files per channel (1048×1048).

### Image acquisition 2D

Similarly, we imaged the 2D plates at 20X (res=0.34µm/pixel) by using Nikon’s perfect focus system (PFS) to capture nine sites per well (2048×2048), 16-bit, in five fluorescent channels (Supplementary table 2).

### Feature extraction and preprocessing intensity projections

CellProfiler was used for the generation of Maximum Intensity Projections. We implemented a quality control based on the PowerLogLogSlope per channel, gave a flag to each image above the 93^th^ percentile and excluded images that were flagged more than once. We then constructed a segmentation which identified each spheroid. From the MIPs, we then extracted object morphologies, intensities and textures. Cell Profiler pipelines are provided. In Jupyter notebooks, we first removed blurry spheroids, and then preprocessed features by standardizing, clipping (±40), and feature selection (blocklisted, variance threshold, and correlation threshold).

### Feature extraction and preprocessing single cells in 3D

We processed and analyzed images with the open-source image analysis software CellProfiler version 4.2.1 (Stirling et al., 2021) and fitted with the Cellpose v2.0 plugin for cellular segmentation (Pachitariu & Stringer, 2022). No illumination correction was performed. In short, we continued training the cyto2 CellPose model to either the nuclei: HOECHST or cells: PHAandWGA images representative of the screen (i.e. different drugs and different sections). For easier labeling, we first used Fiji to subtract the background, actual training and detection was done on raw images. CellProfiler pipelines and the two CellPose models are made available.

In Jupyter notebooks, we preprocessed single cell features by first removing NaNs, aggregated sections using the median, and standardized data both per plate and per optical section. Afterwards, we aggregated well-level features also using the median, followed by clipping them at ±40, and finally feature selection (variance & correlation threshold, dropNA) in Pycytominer (G. Way et al., n.d.).

### Feature extraction and preprocessing single cells in 2D

We processed and analyzed images using the open-source image analysis software CellProfiler version 4.2.5 (Stirling et al., 2021). We ran a quality control pipeline in Cell Profiler 4.0.7, and used an outlier approach to generate flags in Jupyter Notebooks. Illumination was corrected as previously described in (Rietdijk, Tampere, et al., 2022). To segment cells and nuclei, we used the Cellpose v2.0 plugin for cellular segmentation (Pachitariu & Stringer, 2022) and further trained the cyto2 model for each cell line (models provided). We then extracted phenotypic features per cell line (pipelines are provided), followed by Pycytominer for further feature preprocessing (G. Way et al., n.d.). In short, features were first mean-aggregated over sites, then median-aggregated over wells, followed by standardization per plate.

We clipped data at values of 40 and −40. We removed the variance threshold, and correlation threshold.

### Data analysis and visualization

We implemented data analysis in Jupyter notebooks consistently for each data type: MIP, 3D, or 2D. Grit scores, which quantify both the strength and robustness of morphological perturbations (G. Way et al., 2020/2023), were calculated by using the cytominer-eval package (G. Way et al., 2020/2024).

Percent replicating and pairwise technical replicate correlations were determined following the methods described in (Cimini et al., 2022). UMAPs were generated using the UMAP-learn package (McInnes et al., 2020).

Pairwise correlations were visualized using the seaborn cluster map function. Drugs with grit scores below 1.96 were excluded to ensure relevant clustering, and Ward’s linkage method was applied for hierarchical clustering. The cosine similarity was used as distance metric were applicable.

To display orthogonal slices of the spheroids, we made the data isotropic using Clij2 (Haase et al., 2020).

## Data availability

Raw images and cell features will be made available in suitable repositories.

## Code availability

All analysis code is available in GitHub: https://github.com/pharmbio/colopaint3D. SphereDetect is an open-source Python package available https://github.com/Ionshiv/SphereDetect.

## Acknowledgements

We want to thank A. Wenz, J. Rietdijk, and M. Lapins for their kind help in setting up cell culture and analysis, as well as fruitful discussions. We gratefully thank all at Bergman Labora for their help with setting up the Nikon system, and continuous support (M. Andersson, O. Gardner, and G. Lundberg). We used BioRender.com and Adobe Illustrator 2023 for graphical illustrations. ChatGPT has been used as a revision-assistant in writing this manuscript. This project has received funding from the Norwegian Research Council under the International mobility grant agreement no 326003. O.S. acknowledges funding from the Swedish Research Council (grants 2020-03731, 2020-01865, 2024-03566, 2024-04576), FORMAS (grant 2022-00940), Swedish Cancer Foundation (22 2412 Pj 03 H), and Horizon Europe grant agreement #101057014 (PARC) and #101057442 (REMEDI4ALL).

---

**Supplementary Figure 2:**
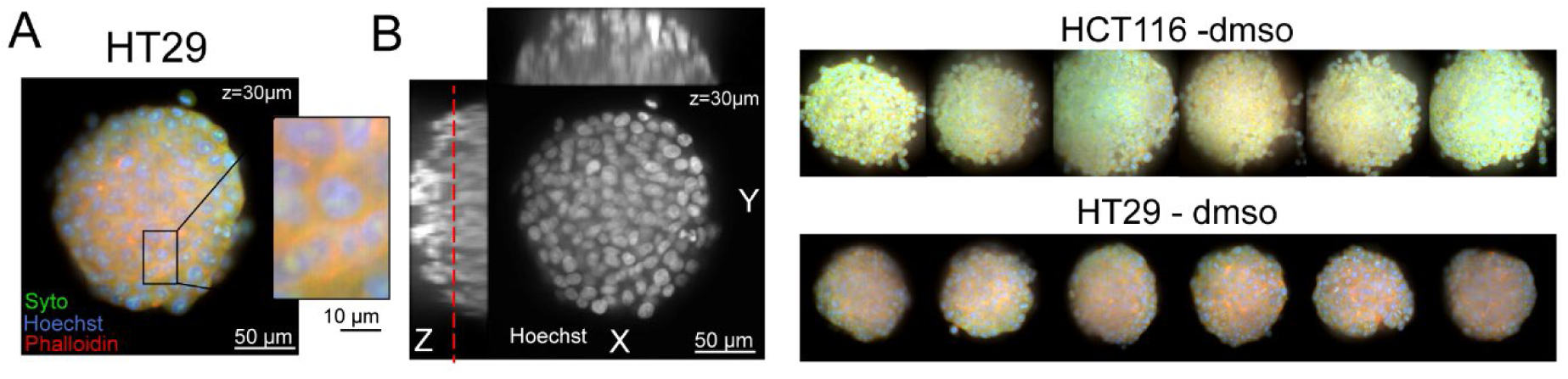
Examples of negative controls. (A) Composite image of a HT29 spheroid at 30µm depth depicting RNA and cytoplasm with (Syto14; green), nuclei with (Hoechst; blue) and cytoskeleton plus Golgi (Phalloidin+WGA; red). Individual cells are seen in the inset. (B) Displayed are orthogonal views of the same spheroid in the Hoechst channel. These views consist of 12 made-isotropic slices (ranging from 0 to 60µm, with 5µm spacing). A dashed red line indicates the position of A (left). (C) Six representative examples of HCT116 (top) and HT29 (bottom) spheroids treated with dmso. Scale bars are 50µm.

**Supplementary Figure 3:**
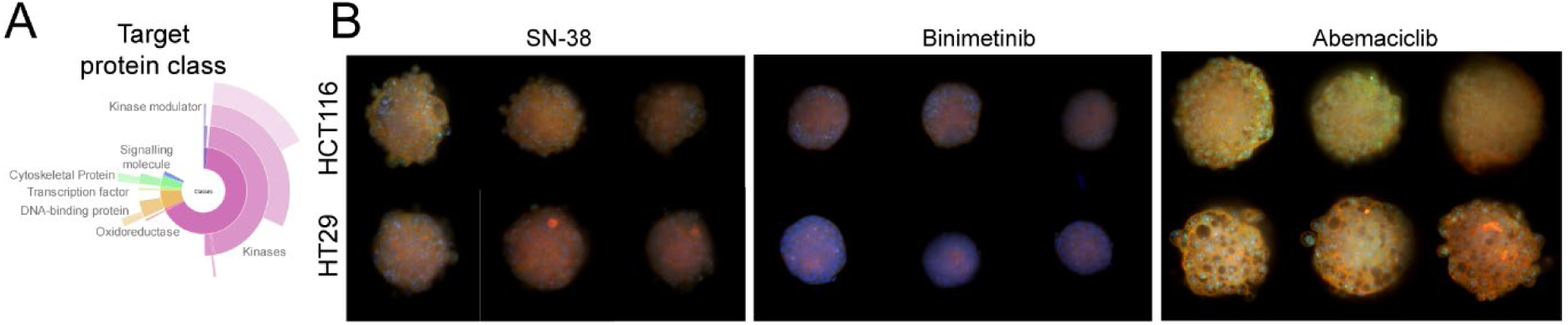
Examples of treated spheroids. (A) A diagram depicting the various targets in our drug library. The figure was created using the clue.io website. (B) Examples of HCT116 (top) and HT29 (bottom) single sections at 30µm depth for the compounds SN-38 (0.03 uM), binimetinib (10 µM), abemaciclib (3.0 uM), and dmso (0.1%). Syto (RNA) is shown in green, Hoechst (nuclei) is shown in blue, and Mitotracker (mitochondria) is shown in red. All scale bars are 50µm.

**Supplementary Figure 4:**
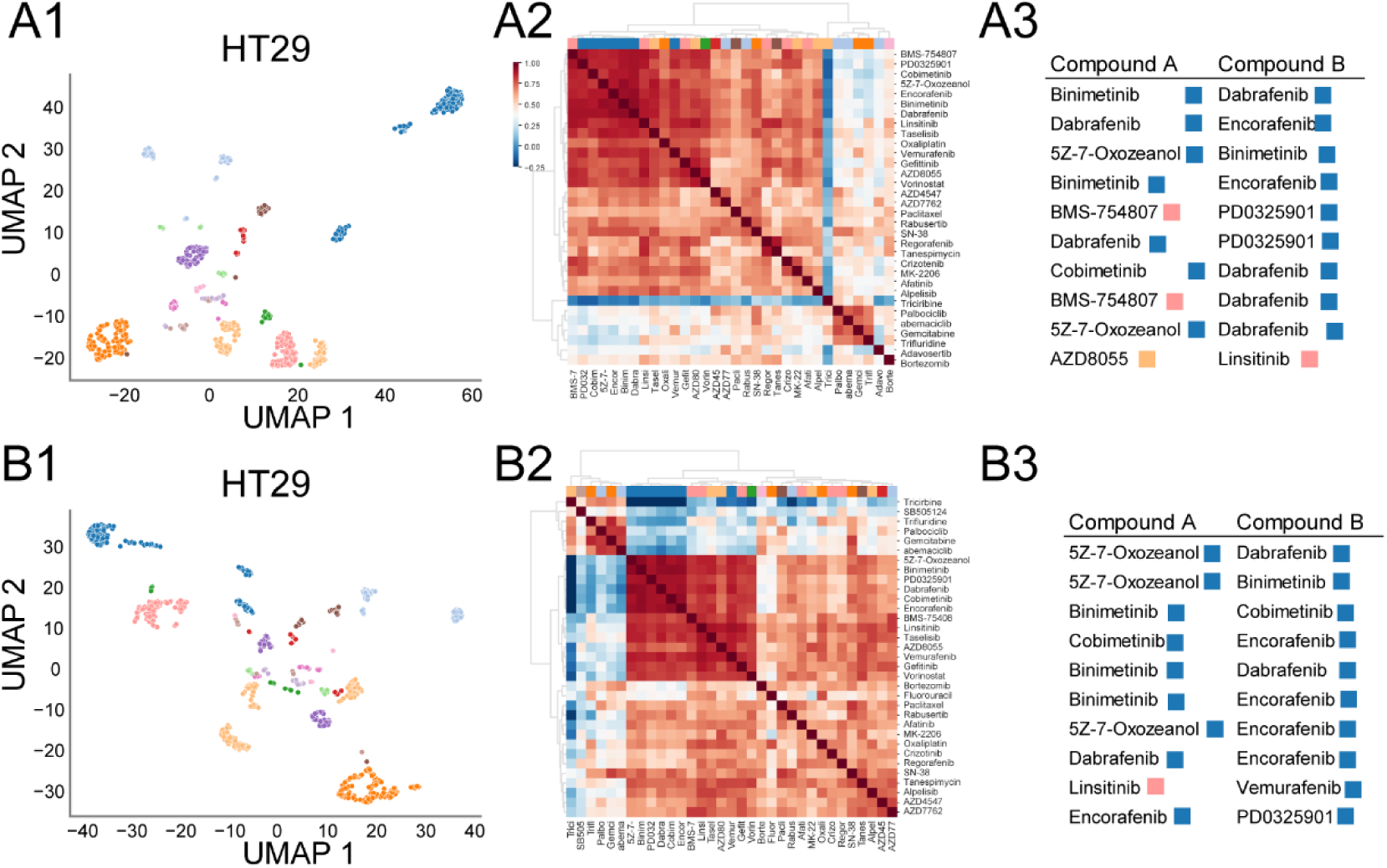
Single-cell morphological profiles facilitate the detection of biologically relevant clusters. (1) Semi-supervised UMAPs of MIP (A1) and aggregated (B1) morphological profiles. (2) Heatmap displaying hierarchical clusters of cosine similarities for MIP (A2) and aggregated (B2) profiles. The ward algorithm was used to create the dendrogram. (3) Top 10 most similar compounds for MIP (A3) and aggregated (B3) profiles. In all plots, pathways are color-coded using the same colormap. All data is based on HT29 spheroids.

**Supplementary Figure 5:**
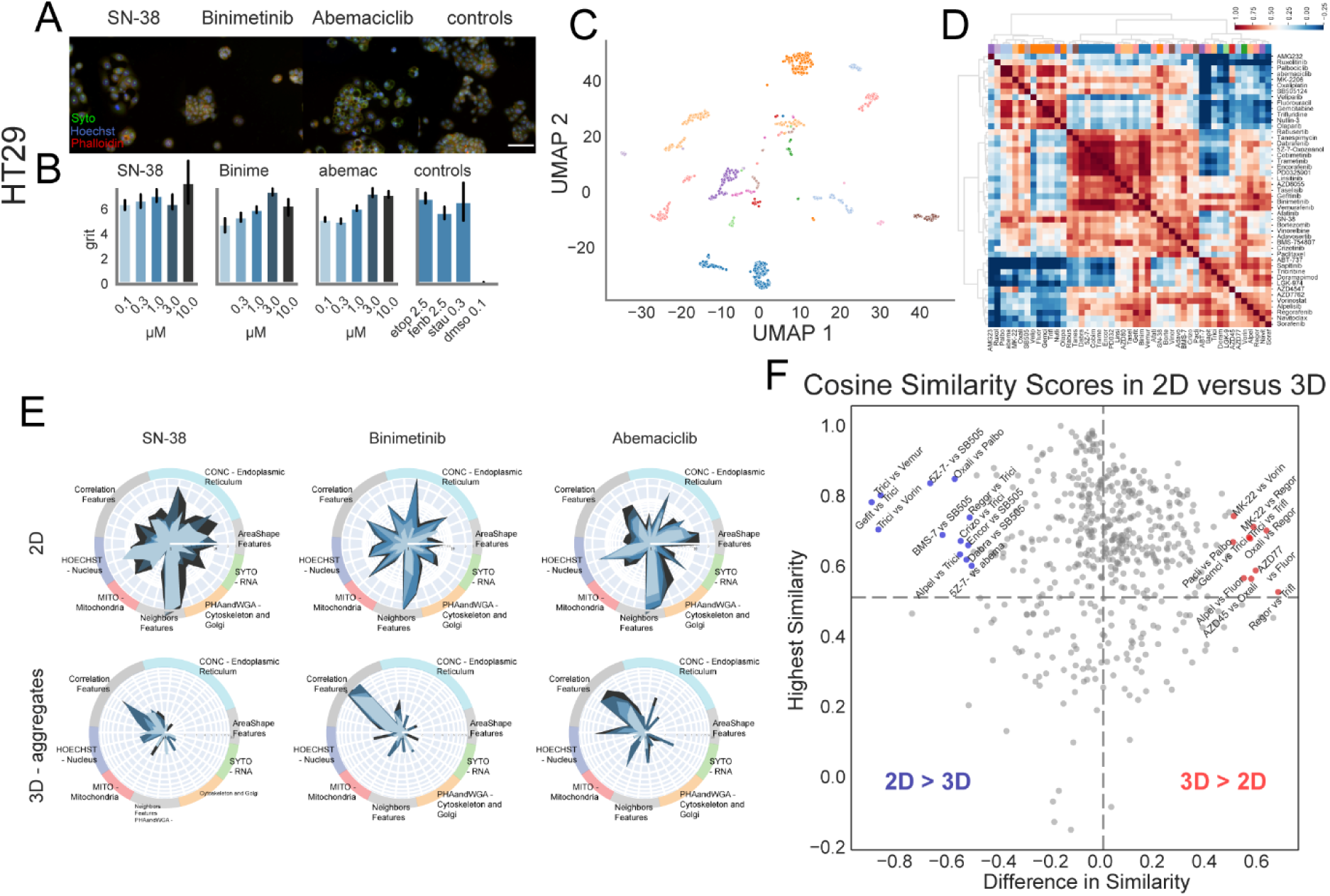
Morphological profiles reveal different responses to DNA damaging compounds in spheroids versus adherent cells. (A) Representative examples of HT29 cultured in 2D for the drugs SN-38 (0.03 uM), binimetinib (10 µM), abemaciclib (3.0 uM), and dmso (0.1%). (B) Grit scores measuring phenotypic effect are shown for 2D aggregate feature profiles. Error bars indicate bootstrapped 95% confidence intervals. (C) Semi-supervised UMAPs of 2D morphological profiles. (D) Heatmap displaying hierarchical clusters of cosine similarities for 2D profiles. (E) Radar plots for the same drugs are displayed for 2D (top) and 3D aggregate (bottom) morphological profiles. Doses are color-coded according to the colors used in B. (F) Comparison of cosine scores between pairs of compounds in 2D and 3D. The x-axis shows the difference in similarity scores (3D – 2D), while the y-axis shows the highest similarity score across both modalities for each pair. We only included pairs with a cosine score of at least 0.5 in either 2D or 3D, and that exceeded the 95% confidence interval cutoff. All data is based on HT29 cells. Scale bar is 250 µm.

